# Comparison of Chemotherapy Effects on Mechanical Sensitivity and Food-Maintained Operant Responding in Male and Female Rats

**DOI:** 10.1101/711358

**Authors:** LP Legakis, CM Diester, EA Townsend, L Karim-Nejad, SS Negus

## Abstract

**Objective:** Chemotherapies of varying classes often cause neuropathy and debilitating chemotherapy-induced neuropathic pain (CINP) sufficient to limit treatment and reduce quality of life for many patients battling cancer. There are currently no effective preventative or alleviative treatments for CINP. Preclinical models have been developed to test candidate CINP treatments; however, studies using these models rarely provide direct comparisons of effects of different chemotherapies or assess the degree to which chemotherapies produce clinically relevant signs of pain-depressed behavior.

**Methods:** Male and female Sprague-Dawley rats received four injections of vehicle, paclitaxel, oxaliplatin, vincristine, or bortezomib on alternate days. Mechanical hypersensitivity, body weight, and food-maintained operant responding were evaluated before, during, and for up to 42 days after initiation of treatment. Morphine potency and effectiveness to reverse chemotherapy-induced effects were also evaluated.

**Results:** All four chemotherapies produced dose-dependent and sustained mechanical hypersensitivity in all rats. Vincristine and oxaliplatin produced transient weight loss and decreases in food-maintained operant responding in all rats, whereas paclitaxel and bortezomib produced lesser or no effect. At four weeks after treatment, operant responding was depressed only in paclitaxel-treated males. Morphine reversed mechanical hypersensitivity in all rats but failed to reverse paclitaxel-induced depression of operant responding in males.

**Conclusions:** Chemotherapy treatments sufficient to produce sustained mechanical hypersensitivity failed to produce sustained or morphine-reversible behavioral depression in rats. Insofar as pain-related behavioral depression is a cardinal sign of CINP in humans, these results challenge the presumption that these chemotherapy-dosing regimens are sufficient to model clinically relevant CINP in rats.

## Introduction

Paclitaxel, vincristine, oxaliplatin, and bortezomib improve the survival and prognosis of patients with many different forms of cancer including breast, ovarian, colon and blood cancers (Crom et al., 1994; de Gramont et al., 2000; Sparano et al., 2008; Bringhen et al., 2010). Despite the effectiveness of these drugs to treat cancers, clinical use of these therapies is often limited by adverse effects that include peripheral neuropathy and associated somatosensory dysfunction (Ramchandren et al., 2009; Ramanathan et al., 2010; Dimopoulos et al., 2011; Seretny et al., 2014; Lee et al., 2017), as well as chemotherapy-induced neuropathic pain (CINP) (Ramanathan et al., 2010; Dimopoulos et al., 2011; Lavoie Smith et al., 2011) and decreased physical functioning (Cersosimo, 2005; Hoffman et al., 2013; Khan et al., 2014; Serrano et al., 2014; Davies et al., 2016; Miaskowski et al., 2017). At present, there are no adequate treatments to prevent or reverse chemotherapy-induced neuropathy, CINP, or pain-related functional impairment (Dworkin et al., 2010; Finnerup et al., 2015).

Administration of chemotherapy to rodents has been used as a noxious stimulus to model chronic pain, neuropathic pain, and CINP. Most commonly, paclitaxel is the chemotherapy tested, and it reliably produces hypersensitive paw-withdrawal responses from mechanical stimuli that can last for weeks to months (Polomano et al., 2001; Pascual et al., 2010; Boyette-Davis et al., 2011; Hwang et al., 2012; Ko et al., 2014; Toma et al., 2017). In rodents, numerous treatments have been identified that alleviate chemotherapy-induced mechanical; however, none of these medications have proven to be effective in clinical treatment of either CINP or neuropathic pain (Xiao et al., 2009; Tatsushima et al., 2011; Paton et al., 2017). Investigating chemotherapy effects on pain-depressed operant behaviors such as in intracranial self-stimulation (ICSS) and food-maintained responding are advantageous for two reasons. First, behavioral depression and functional impairment are cardinal signs of CINP in humans (Dworkin et al., 2010; Mason et al., 2013), and novel preclinical procedures are being developed to evaluate expression of behavioral depression and functional impairment that are reflective of chronic pain (Andrews et al., 2012; Cobos et al., 2012; Toma et al., 2017; Legakis et al., 2018). Second, candidate drugs producing motor impairment can produce analgesic-like but false-positive decreases in assays of hypersensitive withdrawal responses (Xiao et al., 2009; Tatsushima et al., 2011; Paton et al., 2017), but such drugs would not be expected to reverse pain-related behavioral depression (Negus, 2013).

The goal of the present study was to test the hypothesis that chemotherapy regimens sufficient to produce sustained mechanical hypersensitivity in rats would also produce a sustained depression of operant responding maintained by food delivery. Effects were compared for four chemotherapies that have different mechanisms of action to reduce cancercell growth and produce overlapping profiles of side effects that include neuropathy (Seretny et al., 2014; Tsubaki et al., 2018). Morphine, an opioid analgesic used with considerable frequency but marginal effectiveness to treat CINP (Eisenberg et al., 2005; Dworkin et al., 2010), was evaluated for its effectiveness to reverse chemotherapy-induced mechanical hypersensitivity and behavioral depression.

## Methods

### Subjects

Studies were conducted in adult male (65) and female (67) Sprague-Dawley rats (Envigo, Somerset, NJ) with initial weights ranging from 356 to 504 g in males and 234 to 320 g in females. Rats were individually housed and maintained on a 12-h light/dark cycle with lights on from 6:00 AM to 6:00 PM in an AAALAC International-accredited housing facility. Rats in studies of food-maintained operant responding (referred to below as “food-restricted” rats) had access to 45 mg food pellets (BioServ Dustless Precision Pellets, Flemington, NJ) during operant behavior sessions, and they were given access to unlimited water. Daily food rations (Teklad standard diet - 19% protein; Envigo) for these rats were provided one hr after behavioral sessions and were titrated to maintain daily body weights within 5% of the running mean for all individual subjects by sex for at least three days prior to chemotherapy treatment and were given daily throughout the entirety of operant experiments (8.5 ± 0.5 g/day daily food rations in males and 6.0 ± 0.5 g/day daily food rations in females). For all other rats (referred to below as “free-fed” rats), food and water were available ad libitum in the home cage. Animal-use protocols were approved by the Virginia Commonwealth University Institutional Animal Care and Use Committee.

### Drugs

Paclitaxel was obtained as a clinically available 6.0 mg/ml solution (Cardinal Health, Richmond, VA,) and diluted in vehicle (8.3% ethanol, 8.3% Cremophor EL, and 83.4% saline) to a final concentration of 2 mg/ml. Vincristine was obtained as a clinically available 1.0 mg/ml solution (Cardinal Health, Richmond, VA,) and diluted in saline to final concentrations of 0.0625, 0.125, and 0.25 mg/ml. Oxaliplatin was obtained as a clinically available 5.0 mg/ml solution (Cardinal Health, Richmond, VA,) and diluted in saline to final concentrations of 1.25, 2.5, and 5.0 mg/ml. Bortezomib (LC Labs, Woburn, MA) was dissolved in 5.0% DMSO and saline to final concentrations of 0.0625, 0.125, and 0.25 mg/ml. The “vehicle” for all experiments was a composite vehicle composed of the reagents required to dilute or dissolve the various chemotherapies and was composed of 5.0% DMSO, 5.0% glucose, 8.3% ethanol, and 8.3% Cremophor EL in saline (73.4%). All rats were injected intraperitoneally (i.p.) on four alternate days (Days 1, 3, 5, and 7) with vehicle, paclitaxel (2.0 mg/kg), vincristine (0.0625, 0.125, 0.25 mg/kg), oxaliplatin (1.25, 2.5, 5.0 mg/kg), or bortezomib (0.0625, 0.125, 0.25 mg/kg) using an injection volume of 1 ml/kg. This dosing regimen resulted in cumulative doses of 8.0 mg/kg of paclitaxel, 0.25, 0.50, and 1.0 mg/kg of vincristine, 5.0, 10.0, and 20.0 mg/kg of oxaliplatin, and 0.25, 0.50, and 1.0 mg/kg of bortezomib. The dose range for each chemotherapy was determined based on published studies (Amoateng et al., 2015; Fujita et al., 2015; Yamamoto et al., 2015; Legakis et al., 2018) and on preliminary pilot studies that evaluated doses of each chemotherapy that could be administered without lethality using the designated dosing regimen of four injections administered on alternate days. These pilot studies included doses that were at least double those tested for each chemotherapeutic and were deemed too toxic for further study (lethality in at least one rat). Morphine sulfate (National Institute on Drug Abuse Drug Supply Program) was dissolved in sterile water and administered subcutaneously (s.c.) in a volume of 1.0 ml/mg.

### Mechanical sensitivity testing with von Frey filaments in free-fed rats

#### Overview

Effects of treatment with vehicle or with each dose of each chemotherapy were evaluated in 11 separate groups of six rats each (three male and three female). Male and female rats were included in this experimental design to address the National Institutes of Health mandate to include both sexes in preclinical research. Briefly, baseline mechanical sensitivity thresholds were determined on the day before initiation of vehicle, paclitaxel, vincristine, oxaliplatin, or bortezomib treatment, and thresholds were subsequently redetermined on Days 1, 8, 15, 22 and 29 after initiation of chemotherapy treatment. Body weights were recorded daily, and antinociceptive effects of morphine were determined on Day 29 in all groups. The experimental timeline for these studies are shown in Table 1.

**Table 1:** Timeline of experimental events

| **Day** | **Mechanical Sensitivity Experiment** | **Food-Maintained Responding Experiment** |
| --- | --- | --- |
| -2 - 0 | Predrug mechanical sensitivity threshold determination | Predrug total reinforcements determination |
| 1, 3, 5, 7 | Administration of vehicle, paclitaxel, vincristine, oxaliplatin or bortezomib | Administration of vehicle, paclitaxel, vincristine, oxaliplatin or bortezomib |
| 1 - 29 | Weekly mechanical sensitivity threshold testing on days 1, 8, 15, 22, and 29;<br>Daily body weight determination | Daily operant testing;<br>Daily body weight determination |
| 29 | Morphine Testing | 24-hour sucrose preference test |
| 30-31 | Not Applicable | Mechanical sensitivity threshold testing (30);<br>Adjustment to FR1 operant testing (31) |
| 32-42 | Not Applicable | Demand curve determination<br>(FR 1, 3, 10, 18, 32, 56, 100, 180, 320, 560, 1000) |
| 43 | Not Applicable | Adjustment to FR5 operant testing |
| 44-46 | Not Applicable | Saline (44), morphine (45-46) |

#### Testing procedure

On test days, rats were placed on an elevated mesh galvanized steel platform in individual chambers with a hinged lid and allowed to acclimate for at least 20 minutes before exposure to mechanical stimuli. Von Frey filaments (ranging from 0.4 to 15.0 g and increasing in ~0.25 log increments; North Coast Medical, Morgan Hill, CA) were applied to the plantar surface of each hindpaw, and the threshold stimulus to elicit paw withdrawal was determined in log grams using the “up-down” method as previously described (Chaplan et al., 1994; Legakis et al., 2018; Legakis and Negus, 2018). Filament forces greater than 15.0 g were not used because they physically lifted the paw, and as a result, paw movement could not be reliably attributed to a withdrawal response by the subject.

#### Cumulative morphine testing

Following threshold determinations on Day 29, morphine antinociception was evaluated using a cumulative-dosing procedure. Saline and a sequential series of morphine doses were administered s.c. at 60 min intervals. Each dose increased the total, cumulative morphine dose by 0.5 log units (saline, 0.32, 1.0, 3.2, 10 mg/kg), and mechanical sensitivity thresholds were determined 30 min after each injection.

#### Data analysis

For each test condition, body weight data were averaged across rats, and mechanical threshold data were averaged across paws within a rat and then across rats. Effects of each chemotherapy treatment on body weight and mechanical sensitivity were analyzed by two-way ANOVA, with time after initiation of treatment as a within-subjects factor and chemotherapy dose as a between subjects factor. A significant ANOVA was followed by the Holm-Sidak post-hoc test to compare effects of vehicle with effects of each chemotherapy dose at a given time point. All chemotherapies produced sustained mechanical hypersensitivity that reached or remained at peak effect on the last day of testing (Day 29). Accordingly, data from Day 29 were also plotted as dose-effect curves to permit comparisons of potencies and efficacies. (Note: Only one dose of paclitaxel was tested in this study, so the paclitaxel dose-effect curve shows data from a similar previous study that evaluated multiple paclitaxel doses; (Legakis et al., 2018). Chemotherapy potency (ED_0.8_) was determined by linear regression as the dose required to decrease thresholds to 0.8 log g (an effect level achieved by all drugs), and efficacy was defined as the mean threshold observed with the highest dose of each treatment. For all analyses here and below, statistical analysis was conducted using Prism 7.0 (Graphpad Software Inc., San Diego, CA), and the criterion for significance was non-overlapping confidence limits (potency and efficacy comparisons) or p<0.05 (all other analyses).

Morphine antinociceptive effects on Day 29 were expressed as Percent Maximum Possible Effect (%MPE) using the equation: %MPE = [(Test – Daily Baseline) ÷ (Ceiling – Daily Baseline)] x 100, where “Test” was the threshold determined after a morphine dose, “Daily Baseline” was the threshold determined before any injection on Day 29, and “Ceiling” was the maximum force tested (15 g). Morphine effects on chemotherapy-induced mechanical hypersensitivity were analyzed by two-way ANOVA with morphine dose as a within-subjects factor and chemotherapy treatment as a between-subjects factor. A significant ANOVA was followed by the Holm-Sidak post-hoc test. Additionally, morphine ED50 values and 95% confidence limits were determined by linear regression of data from the linear portion of each morphine dose-effect curve.

### Food-maintained operant responding in food-restricted rats

#### Overview

Studies described above identified the highest doses of each chemotherapy that could be safely administered and confirmed that these treatments were sufficient to produce mechanical hypersensitivity (see Results). Subsequent studies evaluated food-maintained operant responding before, during, and after treatment with vehicle or the highest tolerable dose of each chemotherapy (2.0 mg/kg/day paclitaxel; 0.25 mg/kg/day vincristine; 5.0 mg/kg/day oxaliplatin; 0.25 mg/kg/day bortezomib). These five treatments were examined in separate groups of male (N=6 per group) and female (N=6-8 per group) rats. Baseline body weights and rates of food-maintained responding were established before chemotherapy treatment, and both variables were monitored daily for 29 days after initiation of treatment. Additional studies were conducted after Day 29 as described below to assess sucrose preference, mechanical sensitivity, demand curves for food pellets, and morphine effects on paclitaxel-induced depression of food-maintained responding in males. The experimental timeline for these studies are shown in Table 1.

#### Apparatus

Studies were conducted in sound-attenuating boxes containing modular acrylic and metal test chambers (29.2 x 30.5 x 24.1 cm; Med Associates, St Albans, VT). Each chamber contained a response lever, three stimulus lights (red, yellow, and green) centered above the lever, a 2-W house light, and a pellet dispenser that delivered 45 mg sweetened-food pellets (BioServ, Flemington, NJ) to an aperture beside the lever. Different pellets were used for males (Product# F0042, sugar pellets with dextrose and fructose) and females (Product# F0023, sucrose pellets) because the manufacturer discontinued production of Product# F0042 before studies in females were initiated, and the manufacturer recommended Product# F0023 as the most appropriate replacement. Sufficient sugar pellets remained from male studies to permit initial training with these pellets in females before the females were transitioned to sucrose pellets, and during this transition, there was no difference in pellets earned with the two formulations (data not shown). This transition was completed before establishing stable behavioral baselines as described below. Control of stimulus delivery in the operant chamber and collection of data on lever presses and reinforcements earned were accomplished with a computer, interface, and custom software (Med PC-IV, Med Associates).

#### Training and Testing

Onset of the house light signaled the beginning of 30-min behavioral sessions during which lever presses produced delivery of a food pellet under a fixed-ratio (FR) schedule of reinforcement. The FR was gradually increased from FR 1 to FR 5, and after each pellet delivery, there was 0.5-sec time out period during which the lever lights were illuminated and responding had no scheduled consequences. Training continued until the following criteria for stable responding were met for three consecutive days: (1) subjects earned ≥ 75 reinforcements/session, and (2) the number of reinforcements/session on each day varied by ≤ 5% of each individual subject’s running mean of reinforcements. Once responding stabilized under the FR 5 schedule, a 29-day testing protocol began utilizing the same protocol as described for FR5 with training. Thirty-minute operant behavioral sessions were conducted daily (with occasional exceptions on weekends) throughout the 29-day test period, and vehicle or a chemotherapy dose was administered 2 hr before behavioral sessions on Days 1, 3, 5, and 7.

#### Sucrose preference test and mechanical sensitivity test

Following behavioral experiments on Day 29, rats were exposed to a 24-hr, two-bottle choice assay in their home cages. One bottle was filled with water and the other was filled with 2% sucrose dissolved in water. Bottles were weighed before and after the 24-hr session, and the change in weight for each bottle was calculated in grams. Data are expressed as percentage of 2% sucrose choice using the equation: % Sucrose Preference = [Sucrose grams ÷ (Sucrose grams + Water grams)] x 100. On Day 30 after conclusion of the sucrose-preference test, mechanical sensitivity thresholds were determined in all rats as described above.

#### Demand curve testing

On Days 31 and 32, food pellets were made available under an FR 1 schedule as described above. On each subsequent day (Days 33-42) the FR was increased using the following progression: 3, 10, 18, 32, 56, 100, 180, 320, 560, and 1000. Operant sessions lasted 30 min, and the FR progression was incremented daily in each rat until no pellets were earned.

#### Morphine testing

The only evidence for sustained chemotherapy-induced depression of food-maintained responding was obtained in male rats treated with paclitaxel and tested under an FR 5 schedule of reinforcement (see Results). Accordingly, on Day 43, vehicle- and paclitaxel-treated male rats were returned to the original FR 5 schedule for subsequent assessment of morphine effects on paclitaxel-induced depression of food-maintained responding. On Day 44, saline was administered (s.c.) 30 min prior to operant testing, and responding was still depressed in the paclitaxel-treated male rats relative to the vehicle-treated rats. On Days 45 and 46, 1.0 mg/kg morphine and 3.2 mg/kg morphine (s.c.), respectively, were administered 30 min prior to operant testing in vehicle- and paclitaxel-treated male rats. Morphine doses were selected based on morphine potency to reverse chemotherapy-induced mechanical hypersensitivity (see Results).

#### Data analysis

Body weights were evaluated as described above. The primary dependent measure for food-maintained responding was the total number of reinforcements/session. Data from the final three training days prior to initiation of vehicle or chemotherapy treatment were averaged to produce a mean predrug baseline measure of reinforcements/session for each rat. Once vehicle or chemotherapy treatment was initiated, the number of reinforcements/session was determined daily in each rat on Days 1-29 and expressed as a percentage of predrug baseline for each rat using the equation: % Baseline Reinforcements = (Number of Reinforcements on a Test Day ÷ Predrug Baseline Reinforcements) x 100. Changes in food-maintained responding over time were then averaged across rats and analyzed by two-way ANOVA, with time after initiation of treatment as a within-subjects factor and vehicle or chemotherapy treatment as a between-subjects factor. A significant ANOVA was followed by the Holm-Sidak post-hoc test.

For demand-curve analysis of data collected in each group on Days 31-41, when FR values were progressively increased, the number of reinforcements per session was plotted as a function of FR value. These data were fit using a custom-designed GraphPad Prism template (freely available from the Institutes for Behavior Resources, *http://www.ibrinc.org*) with the Exponential Model of Demand (Hursh and Silberberg, 2008) using the equation *Log Q* = *Log Q_0_* + *k* (*e^-a*Q0*C^– 1*), where *Q* represents reinforcers earned, *Q_0_* represents theoretical number of reinforcers earned if the response requirement were zero, *k* represents log10 value of the greatest number of observed reinforcers earned, *e* represents the base of the natural logarithm, *α* represents a free parameter that is adjusted to minimize the difference between predictions of the equation and each demand curve, and *C* represents the response requirement (i.e., FR). Data were included up to the highest FR value at which at least one reinforcer was earned by at least one subject in a given group. The scaling variable *κ* was fixed to a shared value of 2.56, as this value corresponds to the log10 value of the greatest number of reinforcements earned at any response requirement by any single animal. Demand elasticity (α) and free consumption (Q_0_) values were compared between chemotherapies using one-way ANOVA tests to test chemotherapy effects on reinforcements earned with increasing FR values.

Data from the sucrose preference test and mechanical sensitivity testing were also compared across treatments using one-way ANOVA. A significant ANOVA was followed by a Dunnett’s post-hoc test to compare effects of vehicle with effects of each chemotherapy. Morphine effects on chemotherapy-induced mechanical behavioral depression were analyzed by two-way ANOVA with morphine dose as a within-subjects factor and chemotherapy treatment as a between-subjects factor. A significant ANOVA was followed by the Holm-Sidak post-hoc test.

## Results

### Chemotherapy effects in free-fed rats evaluated for mechanical sensitivity

For free-fed rats used in studies of mechanical sensitivity, the mean±SEM baseline body weights were 394.2±6.3 g (males) and 278.4±3.3 g (females) and the mean±SEM baseline mechanical-sensitivity thresholds were 1.17±0.00 log g (males) and 1.16±0.01 g (females). Supplemental Figure 1 shows the time course of changes in body weight during and after repeated treatment with vehicle and each chemotherapy in rats also tested for mechanical sensitivity. Body weights in these free-fed rats treated with paclitaxel or bortezomib did not differ from vehicle-treated controls, whereas vincristine and oxaliplatin dose-dependently decreased body weights relative to vehicle-treated controls.

Figure 1 shows the time course of changes in mechanical-sensitivity thresholds during and after repeated treatment with vehicle and each chemotherapy. As reported previously in a more extensive dose-effect study (Legakis et al., 2018), 2.0 mg/kg paclitaxel produced mechanical hypersensitivity relative to vehicle-treated controls. Similarly, vincristine, oxaliplatin, and bortezomib also produced dose-dependent mechanical hypersensitivity. This hypersensitivity usually emerged by Day 8 after initiation of treatment, and hypersensitivity reached or remained at peak effect on Day 29.

**Figure 1:**
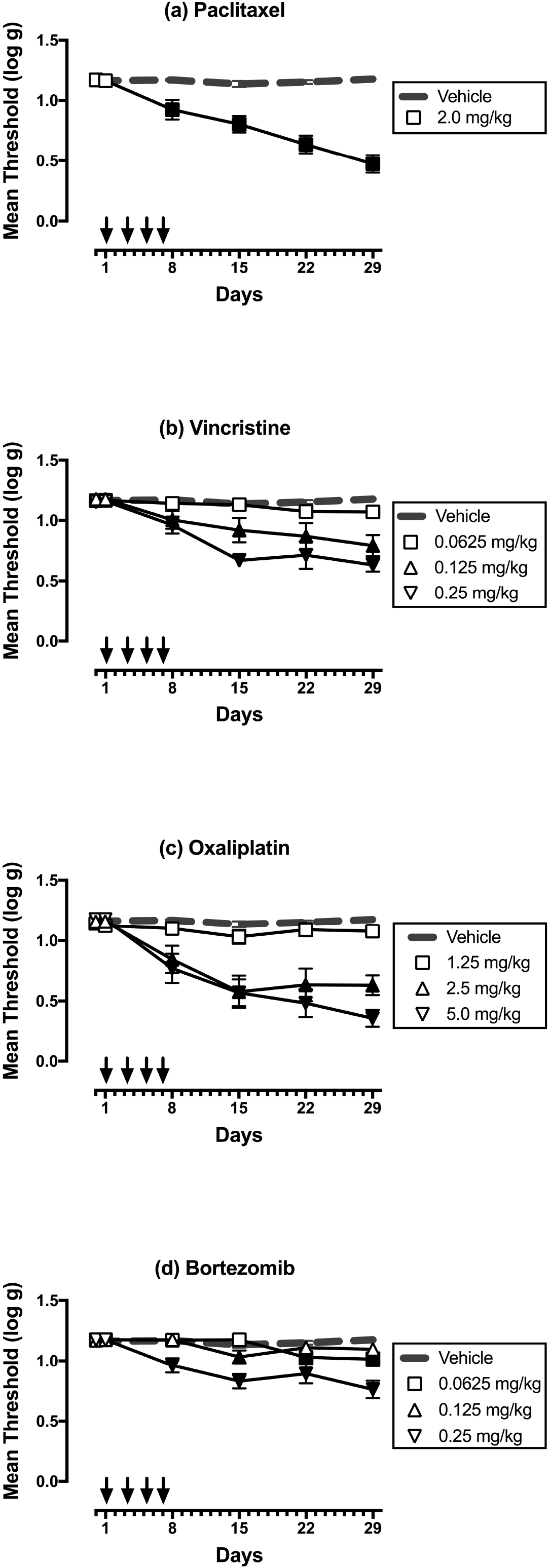
Dose-dependent effects of paclitaxel (a), vincristine (b), oxaliplatin (c), and bortezomib (d) on mechanical sensitivity on Days 1-29. Horizontal axes: Time in days relative to initiation of vehicle or chemotherapy treatment on Day 1. Arrows indicate treatment days. Vertical axes: mechanical sensitivity expressed as threshold stimulation to elicit paw withdrawal in log g. All points show mean±SEM for N=6 rats (3 male and 3 female). Filled points indicate a significant difference from vehicle on a given day as indicated by the Holm-Sidak post-hoc test after a significant two-way ANOVA, p<0.05. Statistical results are as follows. (a) Significant main effects of treatment [F(1,10)=56.15; p<0.0001] and time [F(5,50)=26.74; p<0.0001], and a significant interaction [F(5,50)=27.83; p<0.0001]. (b) Significant main effects of treatment [F(3,20)=14.57; p<0.0001] and time [F(5,100)=23.37; p<0.0001], and a significant interaction [F(15,100)=6.21; p<0.0001]. (c) Significant main effects of treatment [F(3,20)=18.26; p<0.0001] and time [F(5,100)=34.07; p<0.0001], and a significant interaction [F(15,100)=9.99; p<0.0001]. (d) Significant main effects of treatment [F(3,20)=14.98; p<0.0001] and time [F(5,100)=19.28; p<0.0001], and a significant interaction [F(15,100)=7.18; p<0.0001].

Figure 2 compares the dose-effect curves of each chemotherapy to produce mechanical hypersensitivity on Day 29. (Note that paclitaxel data show results for both the present study with 2.0 mg/kg paclitaxel and for a previous study that examined paclitaxel doses of 0.67 and 2.0 mg/kg, Legakis et al. 2018). Potencies and efficacies of each chemotherapy are shown in Table 2. The most potent chemotherapy was vincristine, followed by bortezomib, paclitaxel, and oxaliplatin. The most efficacious chemotherapies were oxaliplatin and paclitaxel, followed by vincristine and bortezomib.

**Figure 2:**
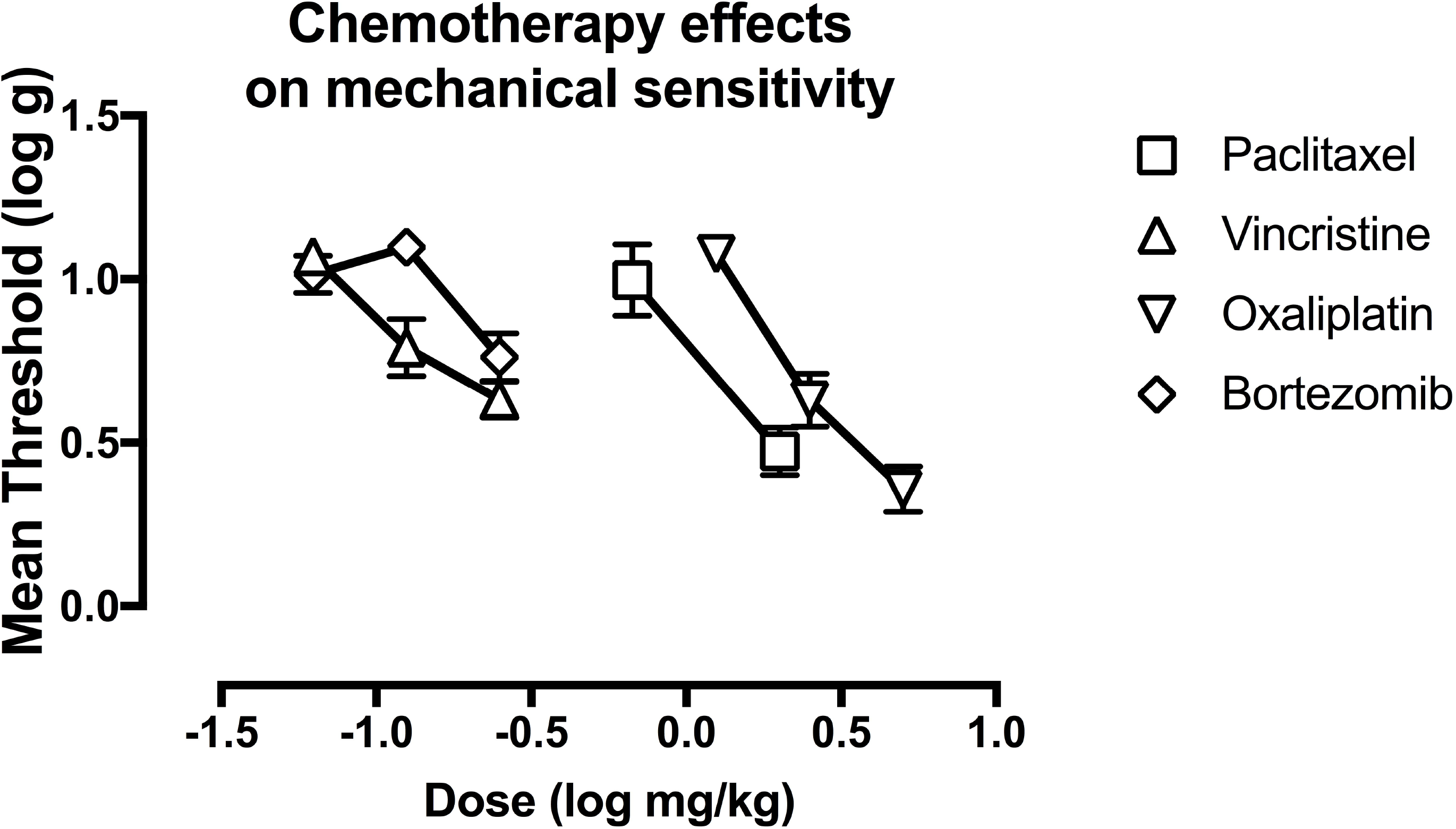
Dose related effects of paclitaxel, vincristine, oxaliplatin, and bortezomib on mechanical sensitivity on Day 29. Horizontal axis: Dose of chemotherapy drug in log mg/kg. Vertical axis: mechanical sensitivity expressed as threshold stimulation to elicit paw withdrawal in log g. All points show mean±SEM for N=6 rats (3 male and 3 female).

**Table 2:** Chemotherapy ED 0.8 and maximum effect values to produce mechanical hypersensitivity.

| Chemotherapy | ED 0.8 (95% CL) in<br>cumulative mg/kg/day | Max Effect (95%CL) in<br>log g |
| --- | --- | --- |
| Paclitaxel | 1.01 (0.65, 1.40) | 0.47 (0.33, 0.61) |
| Vincristine | 0.14 (0.11, 0.19) | 0.63 (0.52, 0.74) |
| Oxaliplatin | 2.01 (1.66, 2.36) | 0.36 (0.22, 0.50) |
| Bortezomib | 0.23 (0.19, 0.34) | 0.76 (0.62, 0.90) |

### Chemotherapy effects in food-restricted rats evaluated for food-maintained responding

For food-restricted rats used in studies of food-maintained responding, the mean±SEM baseline body weights were 338.4±3.2 g (males) and 251.2±2.4 g (females), and the mean±SEM predrug baseline rates of reinforcement per session were 173.9±7.6 (males) and 129.3±4.9 (females). Supplemental Figure 2 shows changes in body weights produced in male and female rats by the maximum tolerable dose of paclitaxel (2.0 mg/kg/day), vincristine (0.25 mg/kg), oxaliplatin (5.0 mg/kg), or bortezomib (0.25 mg/kg). Relative to vehicle controls, body weights in male and female rats were not altered by paclitaxel but were significantly and robustly depressed by vincristine. Oxaliplatin and bortezomib produced smaller decreases in mean body weights in both males and females, but these decreases met criteria for significance only in females.

Figure 3 (a,c,e,g) shows the effects of these same treatments on rates of food-maintained operant responding in males. Relative to vehicle controls, paclitaxel produced small but significant decreases in rates of reinforcement on Days 4, 8, 10, 22, and 24-29. Vincristine, oxaliplatin, and bortezomib each produced significant decreases in rates of reinforcement at various times during the first two weeks of observation, but with these chemotherapies, rates of reinforcement were not different from those in vehicle-treated rats during the last two weeks.

**Figure 3:**
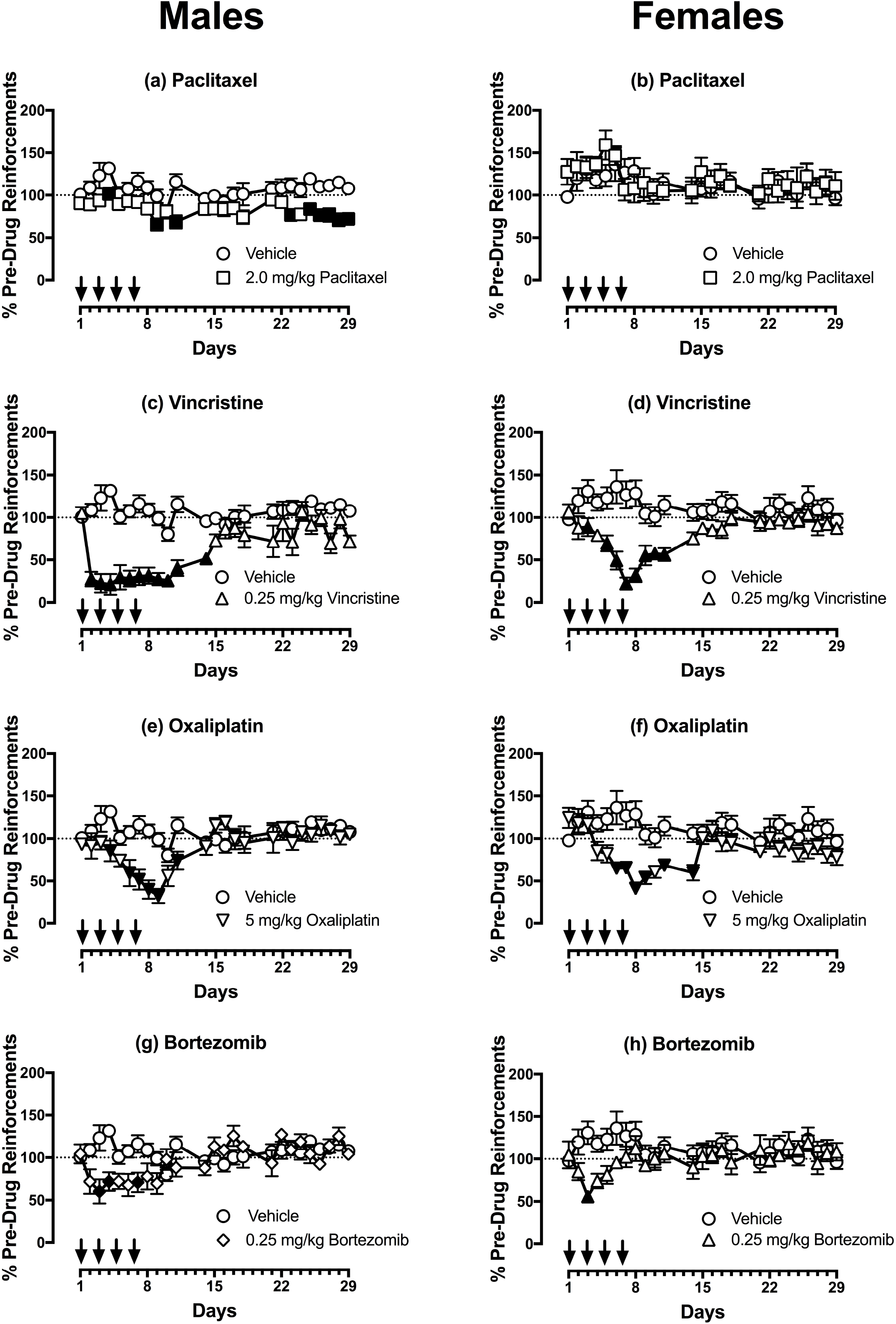
Effects of paclitaxel (a,b), vincristine (c,d), oxaliplatin (e,f), and bortezomib (g,h) on rates of reinforcement in an assay of food-maintained operant responding in males (a,c,e,g) and females (b,d,f,h) on Days 1-29. Horizontal axes: Time in days relative to initiation of vehicle or chemotherapy treatment on Day 1. Arrows indicate treatment days. Vertical axes: reinforcements earned per session expressed as %Pre-Drug Reinforcements per session prior to vehicle/chemotherapy administration. All points show mean±SEM for N=6 male rats and N=6-8 female rats. Filled points indicate a significant difference from vehicle on a given day as indicated by the Holm-Sidak post-hoc test after a significant two-way ANOVA, p<0.05. Statistical results are as follows. (a) Significant main effects of treatment [F(1,10)=23.21; p<0.001] and time [F(24,240)=3.149; p<0.0001], and a significant interaction [F(24,240)=1.90; p=0.008]. (b) No significant main effects of treatment [F(1,11)=0.22; p=0.064] a significant effect of time [F(24,264)=2.02; p=0.004], and a significant interaction [F(24,264)=0.76; p=0.785]. (c) Significant main effects of treatment [F(1,10)=46.31; p<0.0001] and time [F(24,240)=5.98; p<0.0001], and a significant interaction [F(24,240)=6.85; p<0.0001]. (d) Significant main effects of treatment [F(1,13)=31.01; p<0.0001] and time [F(24,312)=2.20; p=0.001], and a significant interaction [F(24,312)=4.50; p<0.0001]. (e) No significant main effect of treatment [F(1,10)=4.95; p=0.050], but a significant effect of time [F(24,240)=6.67; p<0.0001], and a significant interaction [F(24,240)=5.82; p<0.0001]. (f) Significant main effect of treatment [F(1,11)=12.32; p=0.005], a significant effect of time [F(24,264)=2.69; p<0.0001], and a significant interaction [F(24,264)=3.16; p<0.0001]. (g) No significant main effect of treatment [F(1,10)=2.87; p=0.121], but a significant effect of time [F(24,240)=3.12; p<0.0001], and a significant interaction [F(24,240)=4.74; p<0.0001]. (h) No significant main effect of treatment [F(1,11)=2.77; p=0.125], no significant effect of time [F(24,264)=0.96; p=0.520], but a significant interaction [F(24,264)=1.92; p=0.007].

Figure 3 (b,d,f,h) shows the effects each chemotherapy in females. Relative to vehicle controls, paclitaxel produced no change in responding, whereas vincristine, oxaliplatin, and bortezomib each produced significant decreases in rates of reinforcement at various times during the first two weeks of observation; however, rates of reinforcement were not different from those in vehicle-treated rats during the last two weeks.

As an additional assessment of chemotherapy effects on food reinforcement, Figure 4 shows aggregate demand curves for food pellets determined in each group of male and female rats during Days 32-42. Increasing ratio requirements decreased the number of earned pellets in all groups, irrespective of treatment. The exponential model provided a good fit to these data, with mean±SEM R^2^ values of 0.962±0.007 for males and 0.930±0.012 for females (Figure 4c,f). There was no difference across groups in parameters for either free consumption (Q_0_) (Figure 4a,d) or in elasticity of demand with increasing FR value (α) (Figure 4b,e).

**Figure 4:**
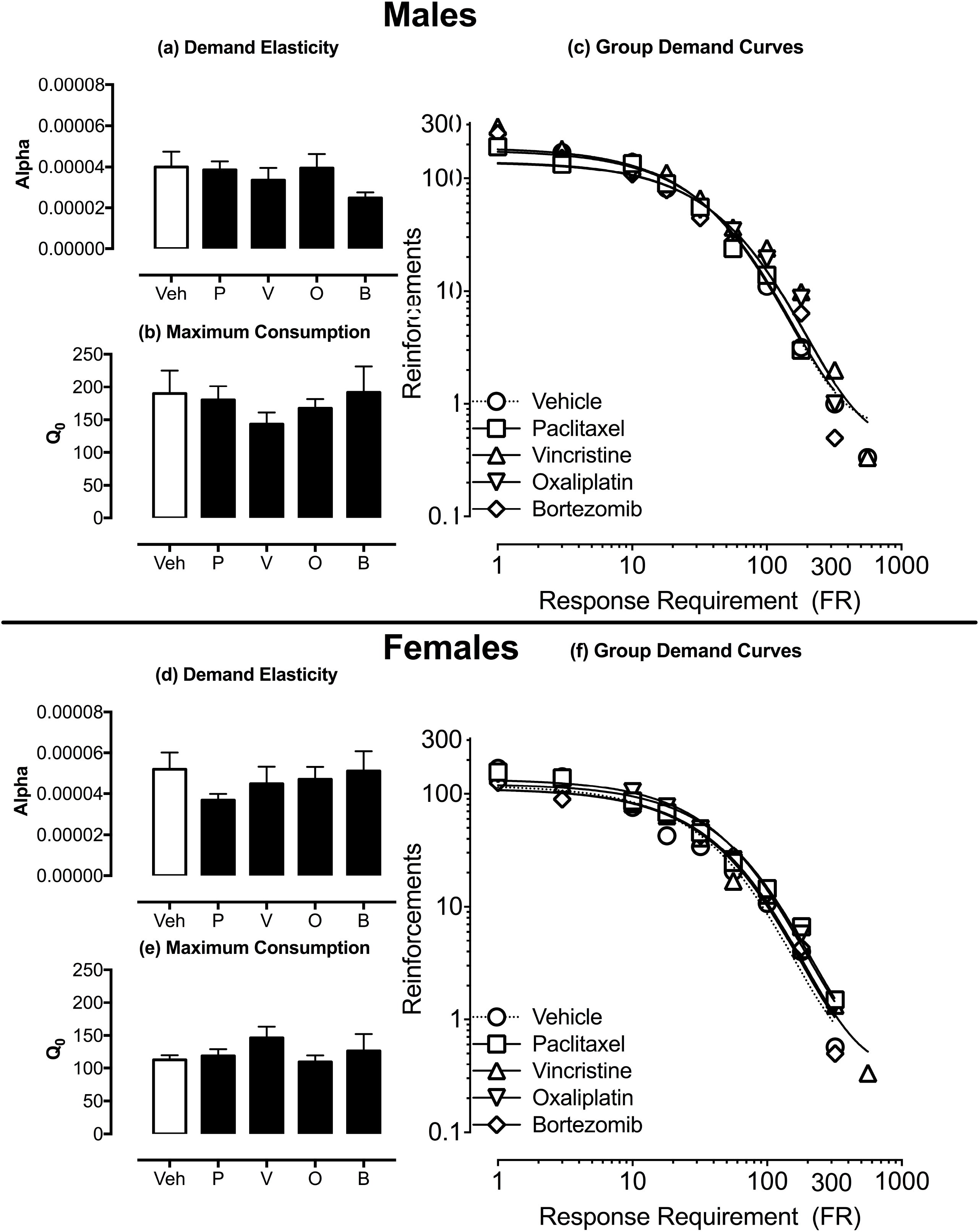
Effects of vehicle or chemotherapy on parameters of demand for food pellets. Panels a, b, d, and e show the effects of vehicle (Veh), paclitaxel (P), vincristine (V), oxaliplatin (O), and bortezomib (B) on demand elasticity alpha (a,d) and maximum consumption Q_0_ (b,e) on Days 32-42. Data for male rats are shown in Panels a-c and female rats are shown in Panels d-f. Panels c and f shows the summary of aggregated essential values across chemotherapies. Statistical results are as follows: (a) No significant effect of chemotherapy treatments [F(4,25)=1.47; p=0.241], (b) No significant effect of chemotherapy treatments [F(4,25)=0.90; p=0.477]. (d) No significant effect of chemotherapy treatments [F(4,25)=1.26; p=0.312], (e) No significant effect of chemotherapy treatments [F(4,25)=0.80; p=0.539].

Figure 5 shows vehicle and chemotherapy effects on sucrose preference (Days 29-30) and mechanical sensitivity (Day 30). Relative to vehicle controls, none of the chemotherapies significantly decreased sucrose preference in either males or females. Paclitaxel and oxaliplatin produced mechanical hypersensitivity in both males and females, consistent with the high efficacy of these chemotherapies to produce mechanical hypersensitivity in the free-fed rats as described above. Vincristine and bortezomib also decreased mean mechanical sensitivity thresholds, but this effect was significant for vincristine only in males and for bortezomib only in females.

**Figure 5:**
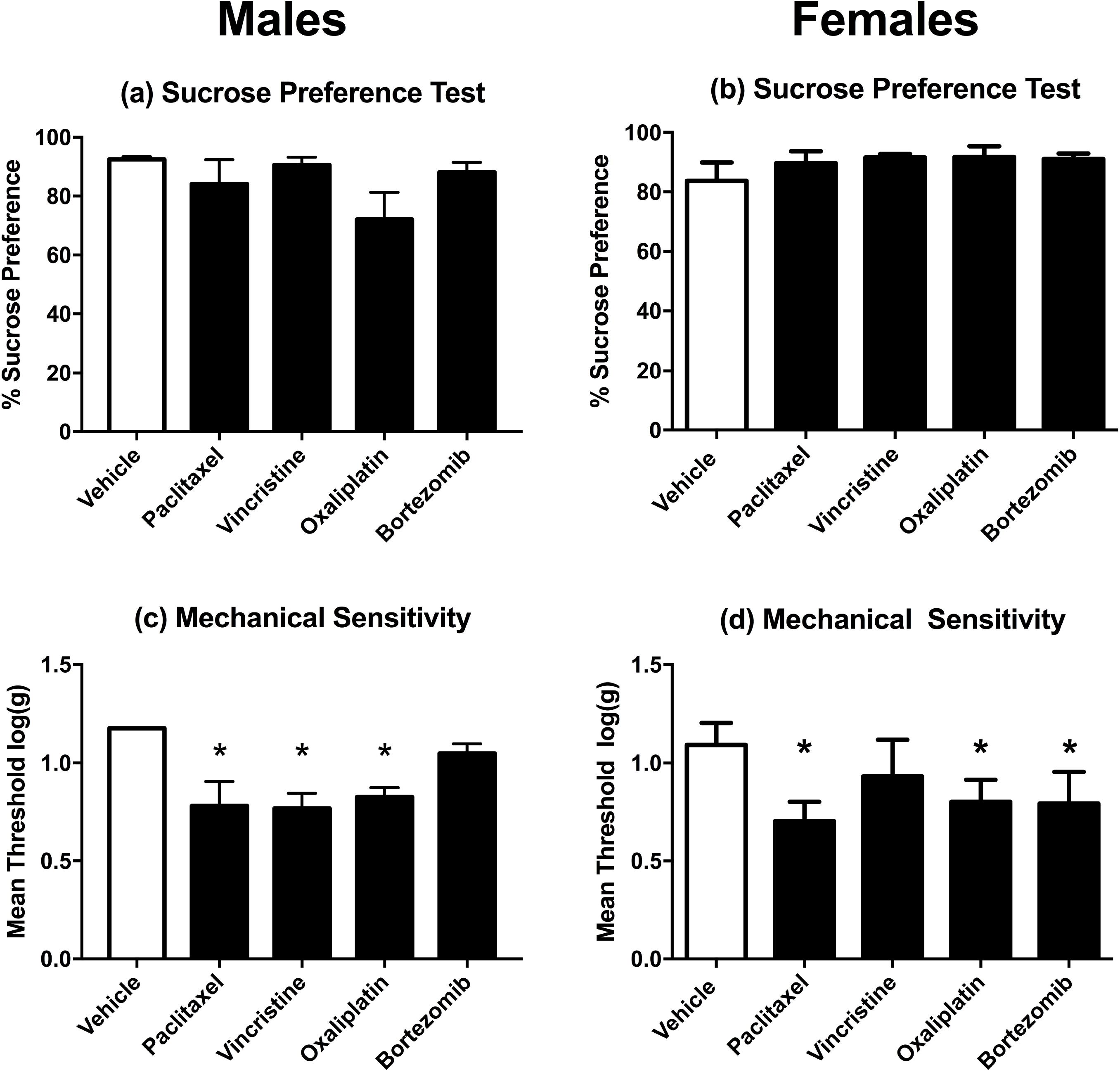
Effects of vehicle or chemotherapy treatment on sucrose preference (a,b) and mechanical sensitivity (c,d) in rats also tested for food-maintained operant responding on Day 29. Data for male rats are shown in Panels a and c and data for female rats are shown in Panels b and d. Horizontal axis: Treatment delivered on Days 1, 3, 5, and 7. Vertical axis: % Sucrose Preference determined on Days 29-30 after treatment initiation (a,b) or mechanical sensitivity expressed as threshold stimulation to elicit paw withdrawal in log g (c,d). All bars show mean±SEM for N=6 male rats and N=6-8 female rats. Asterisks (*) denote significant difference as determined by Dunnett’s post-hoc test after a significant one-way ANOVA, p<0.05 from vehicle. Statistical results are as follows: (a) No significant effect of chemotherapy treatments [F(4,25)=1.93; p=0.138], (b) No significant effect of chemotherapy treatments [F(4,28)=0.82; p=0.516], (c) Significant effect of chemotherapy treatments [F(4,25)=6.46; p=0.001], (d) Significant effect of chemotherapy treatments [F(4,28)=4.88; p=0.004].

### Effects of morphine on chemotherapy-induced mechanical hypersensitivity and behavioral depression

Figure 6a shows morphine effects on mechanical sensitivity thresholds in free-fed rats on Day 29 after treatment with the highest dose of each chemotherapy. Morphine dose-dependently reversed mechanical hypersensitivity produced by all four chemotherapies, and in each case, a dose of 3.2 mg/kg was the lowest dose to produce a significant effect. Table 3 shows that there were no differences across groups in morphine ED50 values to reverse mechanical hypersensitivity.

**Figure 6:**
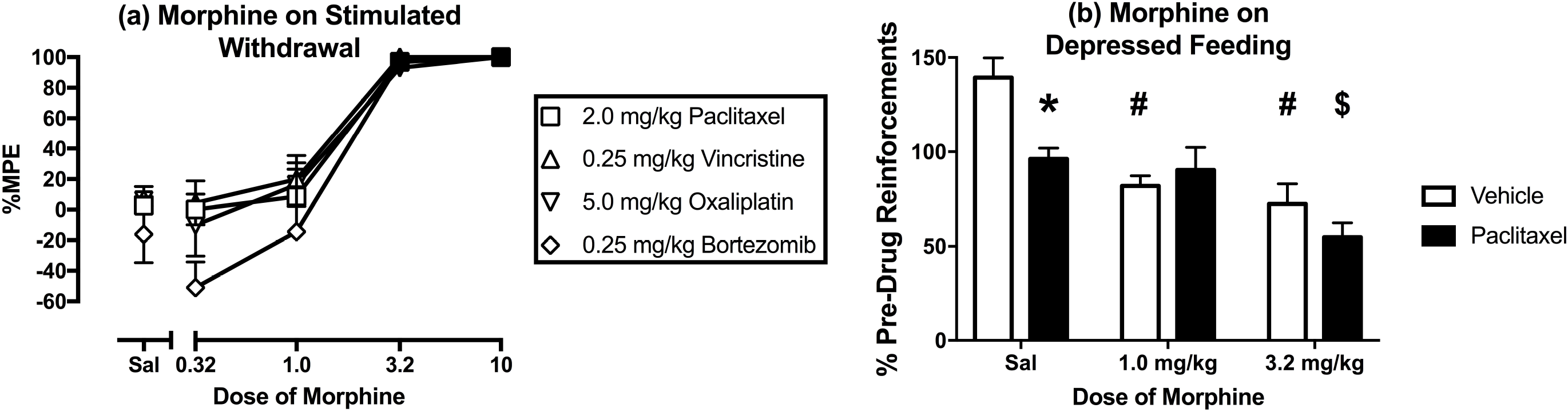
Effects of morphine on mechanical hypersensitivity induced by maximum tolerated doses of paclitaxel, vincristine, oxaliplatin, and bortezomib on Day 29 and on depression of operant responding produced by paclitaxel on Days 44-46. Horizontal axes: Cumulative dose of morphine in mg/kg (log scale) (a) and Dose of morphine in mg/kg (b). Vertical axes: mechanical sensitivity as percent maximal possible effect (%MPE) (a) and reinforcements earned expressed as %Pre-Drug Reinforcements earned prior to vehicle/chemotherapy administration (b). All points show mean±SEM for N=6 rats (3 male and 3 female) in Panel a and mean±SEM for N=6 male rats for Panel b. In Panel a, filled points indicate a significant difference from saline (Sal) at a given dose as indicated by the Holm-Sidak post-hoc test after a significant two-way ANOVA, p<0.05. In Panel b, asterisk (*) indicates significant difference between vehicle- and paclitaxel-treated rats with saline administration, pound (#) indicates significant difference between morphine dose and saline in vehicle-treated rats, and dollar ($) indicates significant difference between morphine dose and saline in paclitaxel-treated rats as indicated by the Holm-Sidak post-hoc test after a significant two-way ANOVA, p<0.05. Statistical results are as follows: (a) No significant main effect of treatment [F(3,20)=1.85; p=0.171], a significant effect of morphine dose [F(4,80)=78.04; p<0.0001], and no significant interaction [F(12,80)=0.757; p=0.692]. (b) Significant main effect of paclitaxel treatment [F(2,20)=5.20; p=0.046], a significant effect of morphine dose [F(2,20)=19.37; p<0.0001], and a significant interaction [F(2,20)=4.37; p=0.027].

**Table 3:** Morphine ED50 values to reverse chemotherapy-induced mechanical hypersensitivity on Day 29.

| **Drug and Dose** | **ED50 in mg/kg**<br>**(95%Confidence Limits)** |
| --- | --- |
| 2.0 mg/kg Paclitaxel | 1.35 (1.01 - 1.83) |
| 0.125 mg/kg Vincristine | 1.03 (0.33 -3.24) |
| 0.25 mg/kg Vincristine | 1.24 (0.60 - 2.56) |
| 2.5 mg/kg Oxaliplatin | 0.85 (0.08, 8.66) |
| 5.0 mg/kg Oxaliplatin | 1.470 (0.44, 4.89) |
| 0.0625 mg/kg Bortezomib | 2.54 (1.52, 4.22) |
| 0.25 mg/kg Bortezomib | 1.01 (0.32- 3.20) |

Figure 6b compares morphine effects on food-maintained responding in vehicle-treated males and in paclitaxel-treated males, the only group of rats that that showed a reduction in operant responding on Day 29 after initiation of chemotherapy treatment. Following data collection for demand curves, the FR value was returned to FR 5, and rates of reinforcement were again significantly lower in paclitaxel-than in vehicle-treated male rats (data not shown). Morphine failed to increase reinforcement rates in paclitaxel-treated male rats. Rather, 1.0 mg/kg morphine significantly decreased reinforcement rate in vehicle-treated rats but not in paclitaxel-treated male rats while also eliminating the differences between groups. The higher dose of 3.2 mg/kg morphine significantly decreased reinforcements earned in both vehicle- and paclitaxel-treated rats, while again eliminating the differences between groups.

## Discussion

The present study compared effects of four mechanistically distinct chemotherapies on mechanical hypersensitivity and motivated behavior in male and female rats. There were three main findings. First, all four chemotherapies produced sustained and morphine-reversible mechanical hypersensitivity across the four weeks of testing, although the chemotherapies did differ in potency and efficacy. Second, chemotherapy efficacy and time course to reduce food-maintained operant responding did not correspond to mechanical hypersensitivity, and in particular, any depression of operant responding usually resolved within a week after termination of chemotherapy administration. Third, in the only exception to this general finding, a sustained depression in food-maintained operant responding was observed in paclitaxel-treated males; however, this paclitaxel effect was not accompanied by a decrease in the reinforcing effectiveness of food as determined by behavioral economic analysis, it was not associated with a decrease in sucrose preference, and it was not alleviated by morphine. Overall, these results suggest that chemotherapy treatments sufficient to produce sustained mechanical hypersensitivity fail to produce evidence of sustained pain-related behavioral depression of positively reinforced operant responding in rats. Insofar as pain-related behavioral depression and functional impairment are cardinal signs of CINP in humans, these results challenge the presumption that these chemotherapy dosing regimens are sufficient to model clinically relevant pain in rats.

### Chemotherapy effects on mechanical sensitivity

The expression of chemotherapy-induced mechanical hypersensitivity in the present study generally agrees with previous studies in rats and mice using paclitaxel (Polomano et al., 2001; Pascual et al., 2010; Boyette-Davis et al., 2011; Hwang et al., 2012; Ko et al., 2014; Toma et al., 2017; Legakis et al., 2018), vincristine (Ji et al., 2013; Linglu et al., 2014; Amoateng et al., 2015), oxaliplatin (Kawashiri et al., 2011; Liu et al., 2013; Fujita et al., 2015), and bortezomib (Chiorazzi et al., 2013; Janes et al., 2013; Yamamoto et al., 2015). However, direct comparison of chemotherapy effects has been complicated by use of different doses and dosing regimens across studies.

To address this issue, the present study used identical dosing regimens with each chemotherapy in free-fed rats to determine full dose-effect curves from ineffective doses up to the highest dose that could be tested without lethality. To facilitate direct comparison, the same dosing regimen was used for all chemotherapies (one injection every other day for a total of four total injections), because this is a commonly used dosing regimen with paclitaxel (Polomano et al., 2001). The effects of these treatments are generally consistent with previous evidence to suggest that chemotherapy-induced mechanical hypersensitivity emerges during the first week of treatment, can be sustained for weeks, and is dose-dependent, with vincristine and bortezomib being more potent than paclitaxel and oxaliplatin (Aley et al., 1996; Polomano et al., 2001; Cavaletti et al., 2007; Ling et al., 2007). To our knowledge, this is the first study to directly compare the efficacies of different chemotherapies to produce mechanical hypersensitivity. A few previous studies have compared effects of single doses of different chemotherapies administered by various treatment regimens, and no clear efficacy differences were evident (Janes et al., 2013; Boehmerle et al., 2014; Hoke and Ray, 2014; Tsubaki et al., 2018). However, the lack of dose-effect determinations in these studies left open the possibility that higher doses could have been tested that might have revealed efficacy differences. Under the conditions of the present study, the order of efficacy to produce mechanical hypersensitivity was (from most to least effective) oxaliplatin, paclitaxel, vincristine, and bortezomib. Only the highest dose of each chemotherapy was administered to food-restricted rats also tested for operant responding; however, these results were also consistent with differences in efficacy insofar as oxaliplatin and paclitaxel produced significant mechanical hypersensitivity in both sexes, whereas vincristine and bortezomib produced significant hypersensitivity in only one sex (vincristine in males and bortezomib in females).

Morphine produced a dose-dependent and complete reversal of mechanical hypersensitivity produced by all four chemotherapies, and the potency of morphine to produce these antinociceptive effects was identical across drugs. This agrees with previous evidence for the potency and effectiveness of morphine to reverse mechanical hypersensitivity induced by paclitaxel, vincristine, and oxaliplatin in rats (Ling et al., 2007; Park et al., 2010; Pascual et al., 2010; Hwang et al., 2012), although morphine sometimes fails to produce a full reversal (Flatters and Bennett, 2004). To our knowledge, this is the first study to report morphine reversal of bortezomib-induced mechanical hypersensitivity.

### Food-Maintained Operant Responding

If chemotherapy-induced mechanical hypersensitivity is a sign of neuropathic pain, then one might also expect concurrent expression of other clinically relevant signs of pain, such as behavioral depression sufficient to interfere with activities of daily living (Dworkin et al., 2008). Moreover, insofar as chemotherapy-induced neuropathic pain manifests primarily as spontaneous shooting, throbbing, or stabbing pain (Kanbayashi et al., 2010; Pachman et al., 2016; Boyette-Davis et al., 2018), one might expect behavioral disruptions in the absence of explicit eliciting stimuli such as the probing with von Frey filaments used to assess mechanical hypersensitivity. To investigate this possibility, the present study examined rates of food-maintained operant responding in chemotherapy-treated rats. Although decreases in operant responding were observed, neither the time course nor the relative effectiveness of chemotherapies to decrease operant responding corresponded to chemotherapy-induced mechanical hypersensitivity. With regard to time course, mechanical hypersensitivity persisted for up to four weeks, whereas any decreases in food-maintained responding usually resolved within a week after termination of chemotherapy treatment (i.e. during the second week). With regard to the magnitude of chemotherapy effects, vincristine produced relatively robust decreases in food-maintained responding while producing relatively weak mechanical hypersensitivity. Conversely, paclitaxel produced relatively small decreases in food-maintained responding while it produced relatively robust mechanical hypersensitivity.

Paclitaxel in male rats was the only treatment to produce a sustained decrease in food-maintained responding that could potentially be related to sustained neuropathic pain. However, even with paclitaxel, this decrease in responding observed under the FR 5 schedule was relatively small compared to decreases produced at earlier times by other drugs, was not accompanied by a significant decrease in the reinforcing efficacy of food as assessed by behavioral economic analysis, was not associated with a decrease in sucrose preference, and was not apparent in females. Thus, the effect was expressed under a very narrow range of conditions. Moreover, paclitaxel-induced depression of food-maintained responding was not reversed by the opioid analgesic morphine. Morphine and other mu opioid receptor agonists have been reported to alleviate decreases in food-maintained operant responding produced by other pain manipulations, such as intraperitoneal acid administration and surgical incision (Martin et al., 2005; Stevenson et al., 2006; Martin et al., 2017). The failure of morphine to reverse paclitaxel-induced decreases in food-maintained responding suggests that this paclitaxel effect may not have been related to pain.

The general failure of these chemotherapy treatments to produce sustained depression of food-maintained responding, despite producing mechanical hypersensitivity, agrees with the failure of paclitaxel to depress operant responding maintained by electrical brain stimulation in rats (Legakis et al., 2018). These findings with chemotherapy-induced neuropathy also agree with the failure of a surgical neuropathy manipulation (spinal nerve ligation) to decrease operant responding maintained either by food or by electrical brain stimulation (Ewan and Martin, 2011; Ewan and Martin, 2014; Okun et al., 2016). Moreover, in agreement with the present failure of any chemotherapy to decrease sucrose preference, paclitaxel treatment sufficient to produce mechanical hypersensitivity also failed to produce a sustained reduction of sucrose preference in mice, and paclitaxel also failed to depress nesting behavior in mice (Toma et al., 2017). Similarly, spinal nerve ligation failed to alter hedonic facial expressions to sucrose solutions delivered to the mouth in rats (Okun et al. 2016). The failure of neuropathy manipulations to alter positively reinforced operant responding cannot be attributed to a general insensitivity of operant procedures to noxious stimuli, because operant responding maintained by either food or electrical brain stimulation can be decreased in an analgesic-reversible manner by more acute noxious stimuli such as intraperitoneal acid administration or surgery (Negus, 2013; Ewan and Martin, 2014; Cone et al., 2018). Taken together, these findings suggest that common models of neuropathic pain in rats are relatively ineffective to produce clinically relevant signs of pain-related behavioral depression and functional impairment, and they are therefore of limited use for preclinical research to investigate mechanisms or treatment of these particular signs of neuropathic pain in humans.

In the only exception to this general trend of negative findings from the present study and the broader literature, one study observed that oxaliplatin produced mechanical hypersensitivity that was concurrent with decreases in milk-reinforced behavior in rats (Ling et al., 2017). In this study, rats had to press their orofacial regions against a ring of von Frey-like filaments to receive the milk reinforcer, and the decreases in responding were interpreted to suggest that responding was reduced by oxaliplatin-induced hypersensitivity to this mechanical stimulus. The effects of oxaliplatin on responding in the absence of the noxious stimulus were not reported, so it is unknown if spontaneous pain might have contributed to the depression of responding in this study; however, this study does raise the possibility that preclinical neuropathy manipulations may be more effective to depress behavior if neuropathy is supplemented by an additional acute provocative stimulus that may function as a punisher.

Although chemotherapy treatments alone appear insufficient to produce pain-related behavioral depression of food-maintained operant responding, the chemotherapy regimens tested here did produce transient decreases in operant responding and parallel transient losses in body weight. These effects agree the clinical adverse outcome profiles noted for paclitaxel (Safran et al., 1997), vincristine (Wagner et al., 2010), oxaliplatin (Chau et al., 2001), and bortezomib (Kane et al., 2007), which notably include transient anorexia, nausea, emesis, and weight loss that resolve quickly after termination of treatment. It is possible that these more transient chemotherapy-induced decreases in food-maintained operant responding and body weight may be a result of the emetic effects that chemotherapeutics produce in the absence of rats’ ability to vomit. Further studies may utilize this premise to model expression of adverse chemotherapy effects on feeding and body weight and test treatments with antiemetic or orexigenic effects.

## Figure Legends

**Supplemental Figure 1:** Effects of paclitaxel (a), vincristine (b), oxaliplatin (c), and bortezomib (d) on body weight of free-feeding rats also tested for mechanical sensitivity on Days 1-29 Horizontal axes: Time in days relative to initiation of vehicle or chemotherapy treatment on Day 1. Arrows indicate treatment days. Vertical axes: body weight expressed as % Baseline Weight determined before initiation of treatment. All points show mean±SEM for N=6 rats (3 male and 3 female). Filled points indicate a significant difference from vehicle on a given day as indicated by the Holm-Sidak post-hoc test after a significant two-way ANOVA, p<0.05. Statistical results are as follows. (a) No significant main effect of treatment [F(1,10)=0.66; p=0.437], a significant effect of time [F(24,240)=1520; p<0.0001], and no significant interaction [F(24,240)=0.31; p=0.999]. (b) Significant main effects of treatment [F(3,20)=6.91; p=0.002] and time [F(22,440)=40.74; p<0.0001] and a significant interaction [F(66,440)=2.75; p<0.0001]. (c) Significant main effects of treatment [F(3,20)=11.13; p<0.001] and time [F(22,440)=36.20; p<0.0001] and a significant interaction [F(66,440)=4.12; p<0.0001]. (d) No significant main effect of treatment [F(3,20)=0.88; p=0.467], a significant effect of time [F(22,440)=46.99; p<0.0001], and no significant interaction [F(66,440)=1.66; p=0.189].

**Supplemental Figure 2:** Dose-dependent effects of paclitaxel (a,b), vincristine (c,d), oxaliplatin (e,f), and bortezomib (g,h) on body weight of food-restricted rats in the assay of food-maintained operant responding in males (a,c,e,g) and females (b,d,f,h) on Days 1-29. Horizontal axes: Time in days relative to initiation of vehicle or chemotherapy treatment on Day 1. Arrows indicate treatment days. Vertical axes: body weight expressed as % Baseline Weight determined before initiation of treatment. All points show mean±SEM for N=6 male rats and N=6-8 female rats. Filled points indicate a significant difference from vehicle on a given day as indicated by the Holm-Sidak post-hoc test after a significant two-way ANOVA, p<0.05. Statistical results are as follows. (a) No significant main effect of treatment [F(1,10)=0.06; p=0.816], a significant effect of time [F(24,240)=3.04; p<0.0001], and no significant interaction [F(24,240)=0.27; p>0.999]. (b) No significant main effect of treatment [F(1,11)=0.65; p=0.437], a significant effect of time [F(24,264)=41.02; p<0.0001], and a significant interaction [F(24,264)=1.59; p=0.048]. (c) Significant main effects of treatment [F(1,10)=47.64; p<0.0001] and time [F(24,440)=17.90; p<0.0001] and a significant interaction [F(24,240)=21.87; p<0.0001]. (d) Significant main effects of treatment [F(1,13)=33.13; p<0.0001] and time [F(24,312)=37.11; p<0.0001] and a significant interaction [F(24,312)=21.39; p<0.0001] (e) No significant main effect of treatment [F(1,10)=3.53; p=0.090] but a significant effect of time [F(24,240)=3.15; p<0.0001] and a significant interaction [F(24,240)=2.70; p<0.0001]. (f) Significant main effect of treatment [F(1,11)=18.83; p=0.001] a significant effect of time [F(24,264)=46.00; p<0.0001] and a significant interaction [F(24,264)=8.92; p<0.0001]. (g) No significant main effect of treatment [F(1,10)=0.31; p=0.590] but a significant effect of time [F(24,240)=4.11; p<0.0001], and a significant interaction [F(24,240)=2.74; p<0.0001]. (h) No significant main effect of treatment [F(1,11)=4.17; p=0.066] a significant effect of time [F(24,264)=49.86; p<0.0001], and a significant interaction [F(24,240)=3.92; p<0.0001].

