## Supplementary figures and images for "Comparison of Chemotherapy Effects on Mechanical Sensitivity and Food-Maintained Operant Responding in Male and Female Rats"

### Supp Fig 1

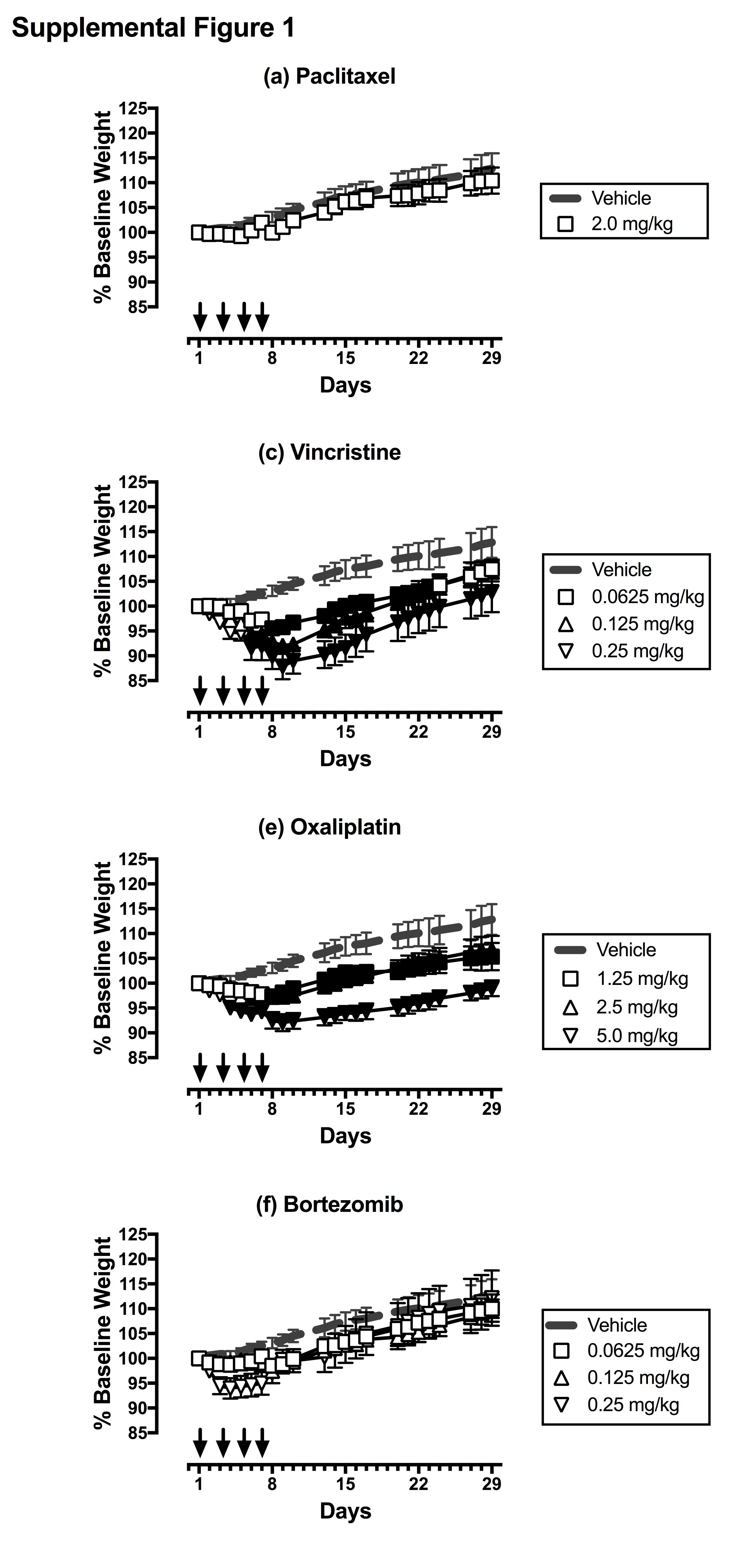

### Supp Fig 2

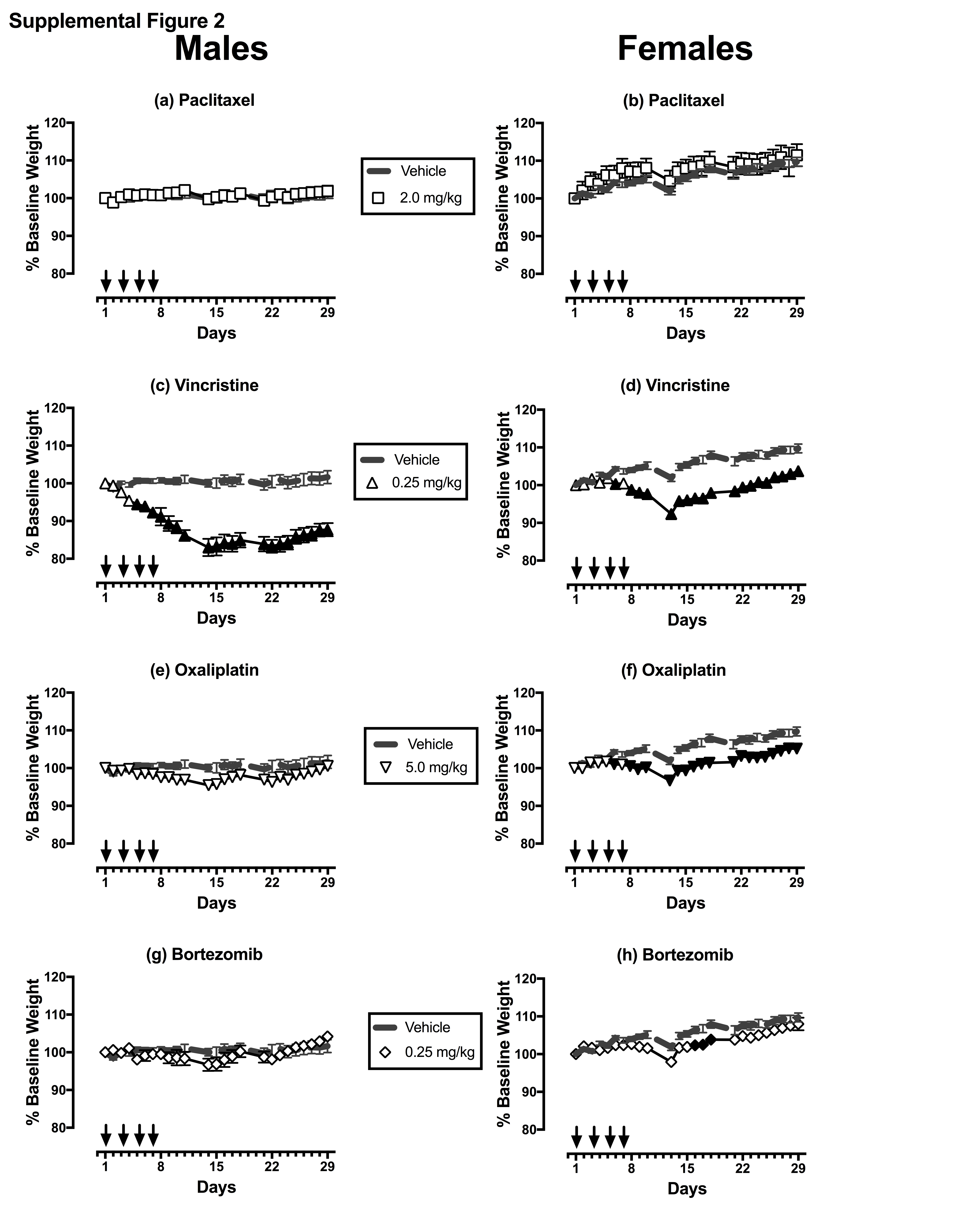
